# Quantification of thermal side effects during pulsed field ablation

**DOI:** 10.1101/2024.11.02.621664

**Authors:** Marcela Mercado-Montoya, Tatiana Gomez-Bustamante, Steven R Mickelsen, Erik Kulstad, Ana González-Suárez, Lawrence J. Overzet

**Affiliations:** IN SILICO STEM S.A.S, Medellín, Colombia; Scripps Health, Electrophysiology, La Jolla, California, United States of America; University of Texas Southwestern Medical Center, Dallas, Texas, United States of America; BioMIT, Department of Electronic Engineering, Universitat Politècnica de València, Spain; University of Texas at Dallas, Electrical and Computer Engineering, Dallas, United States of America

**Keywords:** Pulsed field energy, thermal ablation, atrial fibrillation, computational analysis, in-silico modeling, simulation, numerical analysis

## Abstract

**Background and Aims:** Pulsed field ablation (PFA) has been described as non-thermal, but abundant data exist in oncology applications, and growing data are emerging in cardiology, highlighting that thermal effects are in fact present with PFA. Our objective was to develop a reliable model of the thermal effects arising from PFA of myocardial and esophageal tissue over a range of typical peak voltage operating conditions.

**Methods:** We developed a three-dimensional computer model of the left atrium that can quantify the thermal effects from PFA applications. Energy was applied using a bipolar configuration, and far-field and symmetry boundaries were set as electrically insulating. The model utilizes a monophasic waveform with a pulse width of 100 μs and gap between pulses of 1 s was applied for a total of 50 pulses in a single train, with variations to this easily implemented to accommodate various conditions.

**Results:** Over a range of peak voltage operating conditions (1 kV, 1.5 kV, and 2 kV), minimal temperature rise in the esophagus was seen with 1 kV pulses (corresponding to 215 J input), but with 1.5 kV and 2 kV peak voltages (corresponding to 570 J and 1.23 kJ), temperature elevations reaching 46.3°C and > 62 °C were seen, respectively. These elevations occurred after only a single pulse train of 50 pulses, implying that further elevations in temperature would be seen with subsequent applications. These findings concur with data published in other fields of medicine where pulsed field treatments are utilized.

**Conclusions:** Thermal effects from PFA occur and can be quantified with in-silico modeling. The model described here offers an efficient means of determining temperature effects from PFA over various operating conditions. Energy levels used clinically appear to have the potential to induce collateral thermal injury with repeated applications of pulsed field energy.

## Introduction

Pulsed field ablation (PFA) was originally utilized in cardiology as direct current (DC) ablation in the 1980’s before being supplanted by radiofrequency (RF) ablation as a safer alternative.[1-3] PFA has typically been described as non-thermal, particularly with respect to the predominant mechanism of cell death; however, data from the field of oncology [4-6], as well as growing data in cardiology [7, 8], suggest greater thermal effects from PFA than have been expected.

To further investigate the magnitude of thermal effects during PFA, we developed an in-silico model to serve as a rapid method to quantify thermal effects over various operating conditions without the challenges of animal studies or clinical investigation.

## Methods

We developed a three-dimensional computer model of the left atrium and esophagus to quantify the thermal effects from PFA applications. The model is similar to other published work [9] in which a 3D model represents the PFA procedure with a catheter featuring two cylindrical electrodes (representing momentary anode and cathode) separated by plastic, and assumed to have full contact with the myocardium.

## Geometry

A 3D model representing the PFA procedure (depicted in **Figure 1**) served as the computational framework. This model encompasses a catheter featuring two cylindrical electrodes (representing momentary anode and cathode) separated by plastic. Inspired by the electrode arrangement in PFA basket-like catheters, this design simplifies the configuration to feature just two contiguous electrodes in a horizontal position. Additionally, the model accounts for the myocardium, pericardium (fat layer), and esophagus, each with specific thicknesses (1.5 mm, 0.75 mm, and 2 mm, respectively). The catheter penetrates the tissue by 1 mm.[10] Other tissues are simplified as “other tissues” and share properties similar to the esophagus. Blood flow within the left atrium chamber is also simulated, presumed to move along the x-direction. For computational efficiency, only half of the geometry is considered, exploiting symmetry.

**Figure 1.**
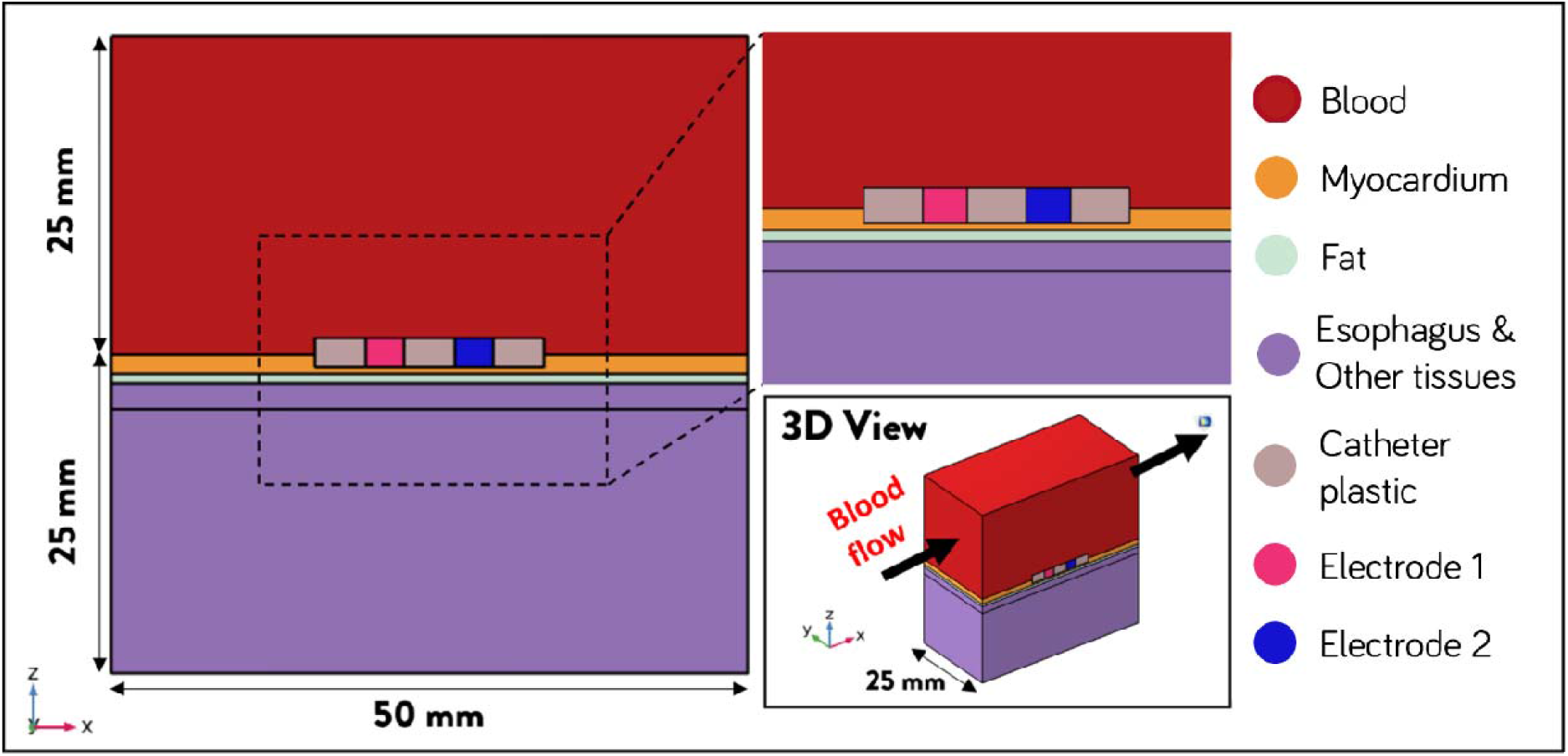
Simplified 3D Model of the Pulsed Field Ablation (PFA) Procedure, illustrating the catheter with two cylindrical electrodes (anode and cathode) and surrounding tissue layers, including myocardium pericardium (fat layer), esophagus, and other tissues. Arrows depict the direction of blood flow within the left atrial chamber.

### Governing Equations and study type settings

The electromagnetic heating and blood flow phenomena involved in the PFA therapy applied in the left atrium endocardial wall was modeled by coupling the electromagnetic equations with the bioheat transfer equation by means of a heat source obtained from electromagnetic power losses and a perfusion heat source from blood in the tissues. The blood flow in the left atrium was modeled through the continuity and Navier-Stokes equations. A stationary study is used for the blood fluid flow, while the time-dependent approach is used for solving electromagnetic heating.

### Boundary and Initial Conditions

Three peak voltage operating conditions in the bipolar configuration were simulated (1 kV, 1.5 kV, and 2 kV). The energy is applied between the two metal electrodes in this configuration with one electrode acting as the positively biased anode and the other as the grounded cathode. An advantage of this configuration is that no dispersive electrode need be considered. Far-field and symmetry boundaries were set as electrically insulating. A monophasic waveform with a pulse width of 100 µs and gap between pulses of 1 s (**Figure 2**) was applied for a total of 50 pulses in a single train.

**Figure 2.**
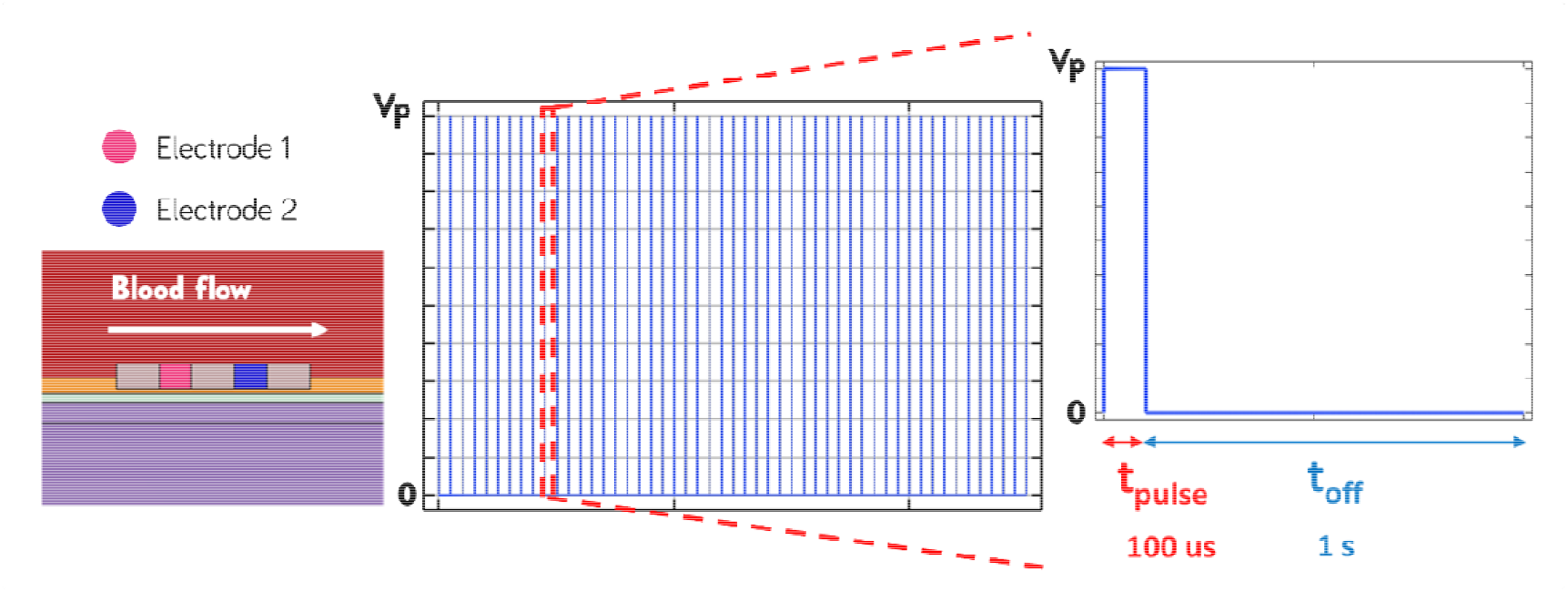
Electrical boundary conditions represented by a bipolar monophasic PFA protocol. The protocol consists of a single train of 50 pulses, each lasting 100 µs, with a 1 s pause between pulses, applied between the anode and cathode.

Blood flow in the model was assumed to be steady-state and directed in the positive x-direction, as illustrated in **Figures 1 and 2**. At the inlet, a normal inflow velocity of 3 cm/s at 37 ºC is prescribed, while a zero-pressure condition is enforced at the outlet. The chosen inlet velocity is approximately 5 times lower than the average value expected in the left atrium for AF patients, which has been reported between 15 and 19 cm/s.[11] However, previous studies have shown that variations in blood flow velocity do not significantly impact the peak temperature results. [12] Non-slip boundary conditions are applied to the walls. Far-field boundaries are set to the body temperature (37 ºC), and symmetry boundaries enforce a zero-flux condition. These boundary conditions are carefully defined to accurately represent the physiological conditions and ensure consistency in the simulation setup.

The time-domain equations, relevant for this study are presented in Equations 1 to 3. Where ***J*** is current density (A/m^2^), is the electric flux density (C/m^2^), is the electric field strength (V/m), *Φ* is electric potential (V), is the permittivity of the material and *σ* is electric conductivity (S/m). The electrical problem is computed in all the subdomains except in the catheter electrodes, as they are modeled as a perfect electric conductor.

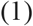

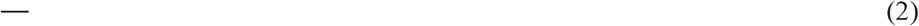

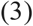

The thermal problem was solved for all the domains using the Bioheat equation (Equation 4).

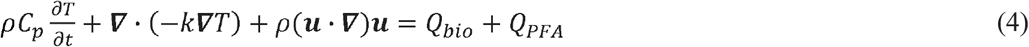

where *T* is the temperature (K = ºC+273.16), *ρ* is density (kg/m^3^), *c*_*p*_ is specific heat capacity at constant pressure (J/kg·K), *k* is thermal conductivity (W/(m^2^·K)) and **u** is the velocity field (m/s) for fluidic subdomains. The thermal advection term is considered exclusively for the subdomains corresponding to the blood and the cold circulating water (i.e. inside the silicone tube).

The heat sources *Q*_*bio*_ and *Q*_*PFA*_ (W/m^3^) correspond to biological and electromagnetic phenomena, respectively. The term *Q*_*bio*_ is composed of metabolic (*Q*_*m*_) and perfusion (*Q*_*p*_): *Q*_*bio*_ = *Q*_*m*_ + *Q*_*p*_. The metabolic heat is negligible compared to energy dissipation and was hence ignored.[13] The perfusion heat was given by Equation 9.

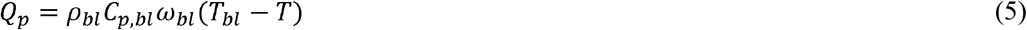

where the following blood properties are involved: *ρ*_*bl*_ is density (kg/m^3^), *C*_*p,bl*_ is heat capacity in (J/kg·K), *ω*_*bl*_ is perfusion rate (s^−1^) and *T*_*bl*_ is blood temperature (310.16 K = 37 ºC). The perfusion rate for each tissue is taken from the ITIS database.[14] The RF-induced heat source is due to Joule effect and is given by Equation 6.

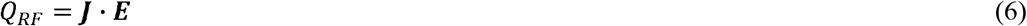

The blood flow was considered incompressible by solving the continuity and Navier-Stokes equations (Equations 7 and 8), and the calculated velocity profile is then coupled to the advection term in the heat transfer equation. In Equations 7 and 8, *ρ* is the fluid density (kg/m^3^), **u** is the velocity field (m/s), µ is the dynamic viscosity (Pa·s), p is the pressure and **f** are the volumetric forces (not considered in this study).

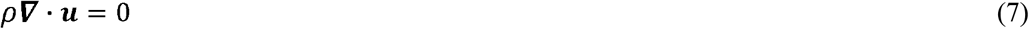

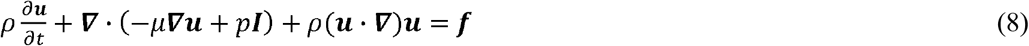

Equations 10 to 12 present the temperature dependence of the heat capacity, thermal conductivity, and density respectively, which are taken from predefined COMSOL functions for tissues, and the reference values from ITIS database.[14]

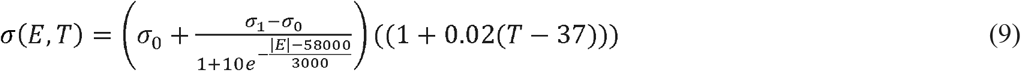

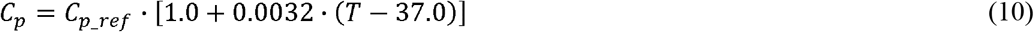

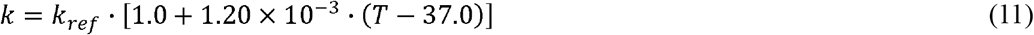

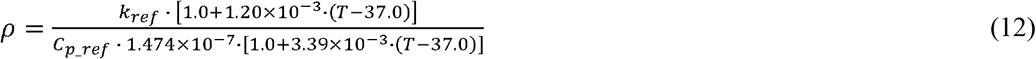

## Materials

The electric and thermal properties of the subdomains are presented in Table 1. Additionally, for the blood domain, dynamic viscosity is also defined, which is useful for solving the Navier-Stokes equations. The irreversible electroporation of tissues is included in the model by defining the electrical conductivity as a function of both electric field and temperature. A sigmoid function dependency of electrical conductivity on electric field describes a tissue with a base electrical conductivity σ_0_ that irreversibly reaches the electroporation conductivity value σ_1_ in the regions where the electric field is greater than 580 V/cm.[15, 16] The values for σ_0_ and σ_1_ are presented in Table 1 and obtained from ITIS database at 10 Hz and 500 kHz respectively.[14] An increase of 2%/ºC was also considered in the electrical conductivity.[17]

**Table 1.**
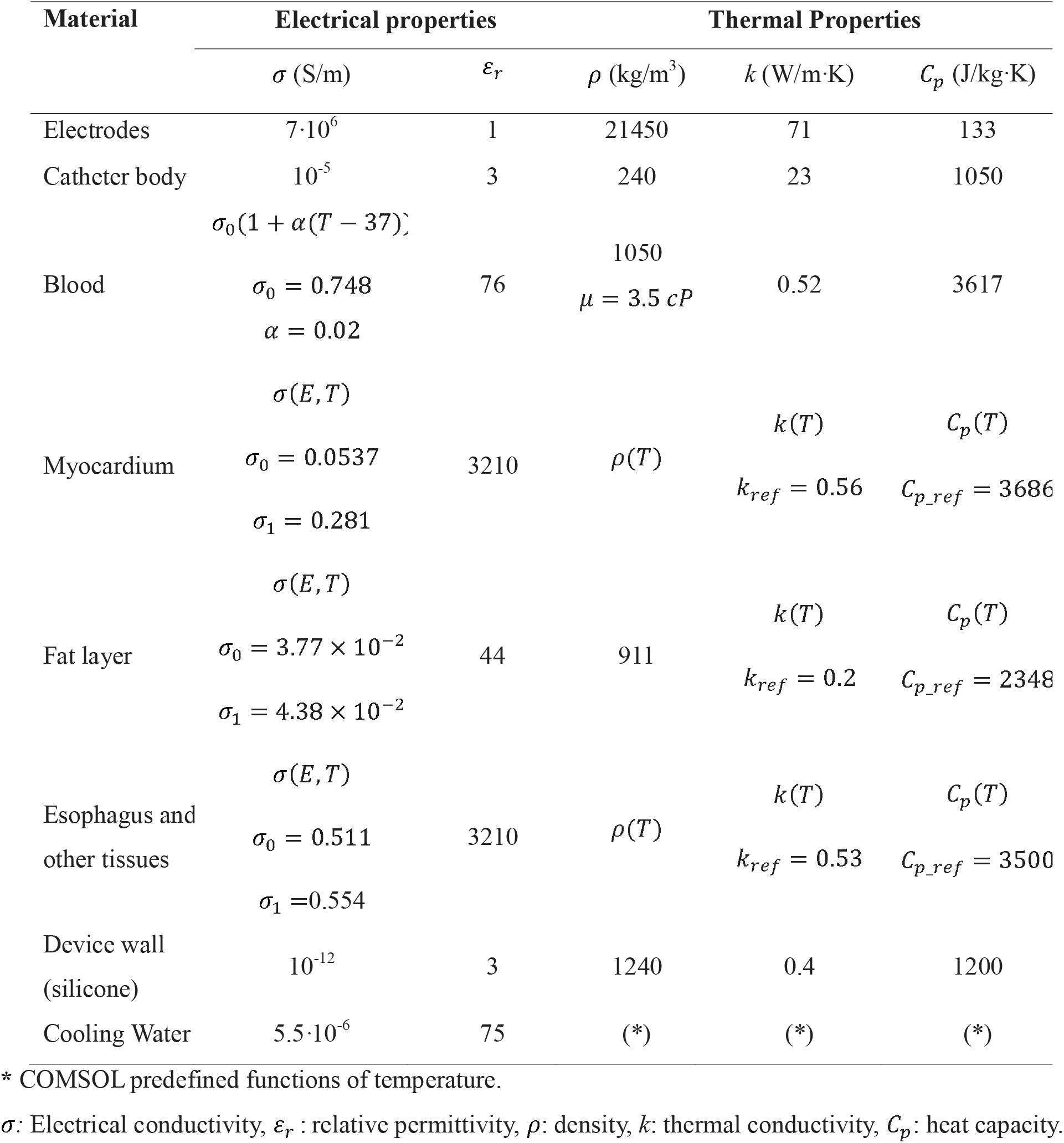
Material properties of the computer model.[14, 15].

### Model Verification

The computational model verification was conducted considering three pulse amplitudes: 1000 V, 1500 V, and 2000 V. The verification process focused on mesh, dimensions, and solver independence. Peak values for temperature and electric field strength in both the myocardium and esophagus during the first and fifth pulses were utilized as convergence criteria. These values were selected based on thermal and electrical analysis. The convergence criteria were defined as follows: the relative error of these variables remains within 5%, and they maintain reasonable values independently of further adjustments in dimensions, solver, and mesh configurations.

### Mesh Independence

The mesh independence results for pulse peak voltage equal to 1000 V, are presented in **Figure 3**. The red line, showing the relative error between Mesh 1 and Mesh 2 (Error 1-2) shows the independence criteria is not fulfilled in the myocardium, neither for the first pulse, nor for the fifth pulse (**Figure 3a**), as its value is over 5%. In contrast, the relative error between Mesh 2 and Mesh 3 (Error 2-3) fulfills the criteria for all the cases in both the myocardium and the esophagus. For the case of maximum electric field, both Error 1-2 and Error 2-3 fulfills the criteria. Based on the previous analysis, the Mesh 2 is the mesh with the minimum number of elements that fulfills the convergence criteria. Nevertheless, as this mesh independence analysis was made considering the minimum voltage considered in this study, which is 1000 V. The Mesh 3 was chosen as the converged mesh.

**Figure 3.**
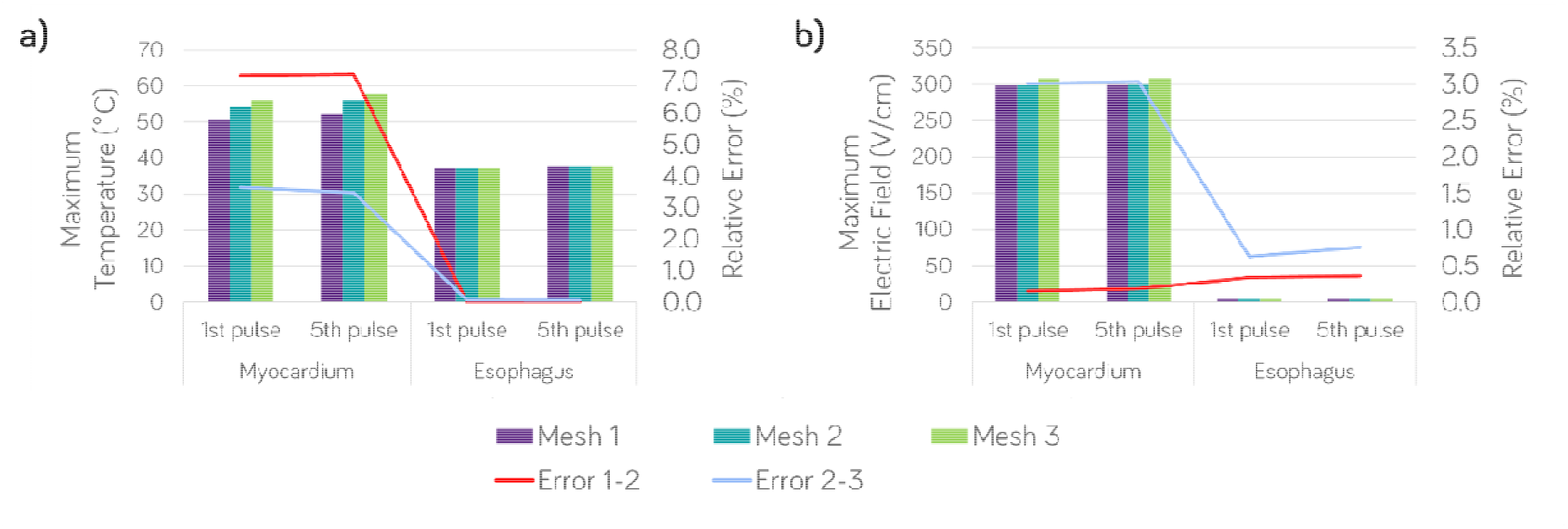
Mesh independence for 1.0 kV, comparing a) maximum temperature and b) maximum electric field in both the myocardium and esophagus at the end of the first and fifth pulse for three different mesh configurations with an increasing number of elements. The relative errors for temperature and electric field between Mesh 1 and 2 (Error 1-2), and Mesh 2 and 3 (Error 2-3), are presented as lines and read on the right y-axis.

### Dimension independence

The results for dimension independence considering 1000 V and 1500 V pulse peak voltages are shown in **Figure 4** and **Figure 5** respectively. As presented in **Figure 4**, both the Error 1-2 and Error 2-3 fulfill the convergence criteria. So, for 1000V, convergence was at 5 cm width when including the infinite elements domain feature. On the other side, based on the results in **Figure 5**, a width of 5 cm with infinite elements is not enough for 1500 V, as the model crashes with no converging results. This problem was solved by adjusting the width to 10 cm and 15 cm. The results for Error 2-3 confirm that 10 cm with the inclusion of infinite element domain feature from COMSOL was enough to fulfill the dimensions independence criteria.

**Figure 4.**
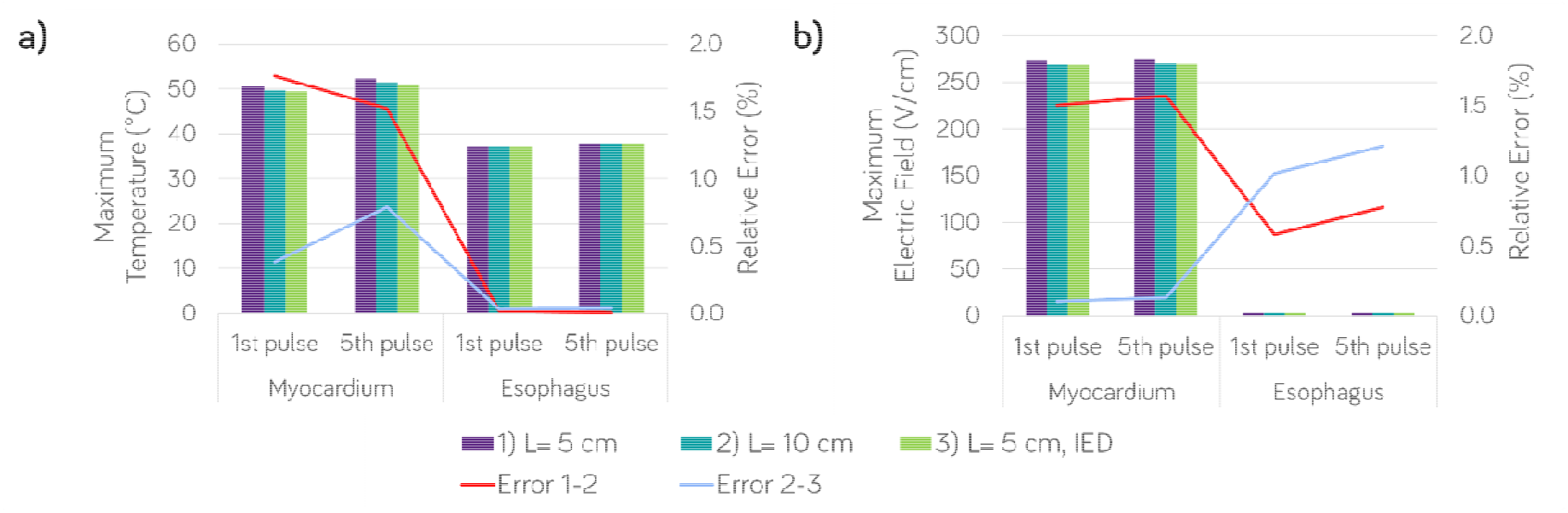
Dimensions independence for 1.0 kV, comparing a) maximum temperature and b) maximum electric field in both the myocardium and esophagus at the end of the first and fifth pulse for three different edge sizes with and without using Infinite Element Domains (IED) feature from COMSOL. The relative errors for temperature and electric field between case 1 and 2 (Error 1-2), and case 2 and 3 (Error 2-3), are presented as lines and read on the right y-axis.

**Figure 5.**
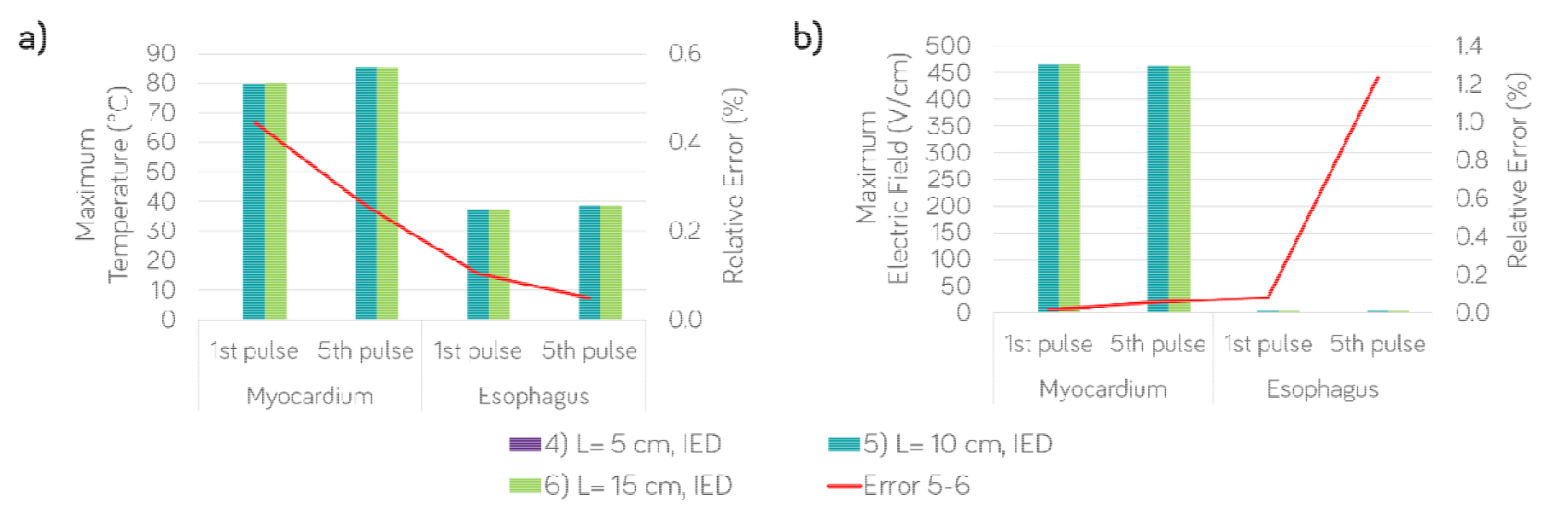
Dimensions independence for 1.5 kV, comparing a) maximum temperature and b) maximum electric field in both the myocardium and esophagus at the end of the first and fifth pulse for three different edge sizes using the Infinite Element Domain (IED) feature from COMSOL. The case 4) did not converge, so the relative errors for temperature and electric field between case 5 and 6, are presented as a red line and read on the right y-axis.

Based on the mesh and dimensions independence studies deployed above, the **Figure 6** presents the final mesh and geometry used for all the peak voltages per pulse considered in this study: 1000 V, 1500 V, 2000V. The mesh evidences the refinement around the electrodes area and the proper sweep type mesh in the infinite elements domain.

**Figure 6.**
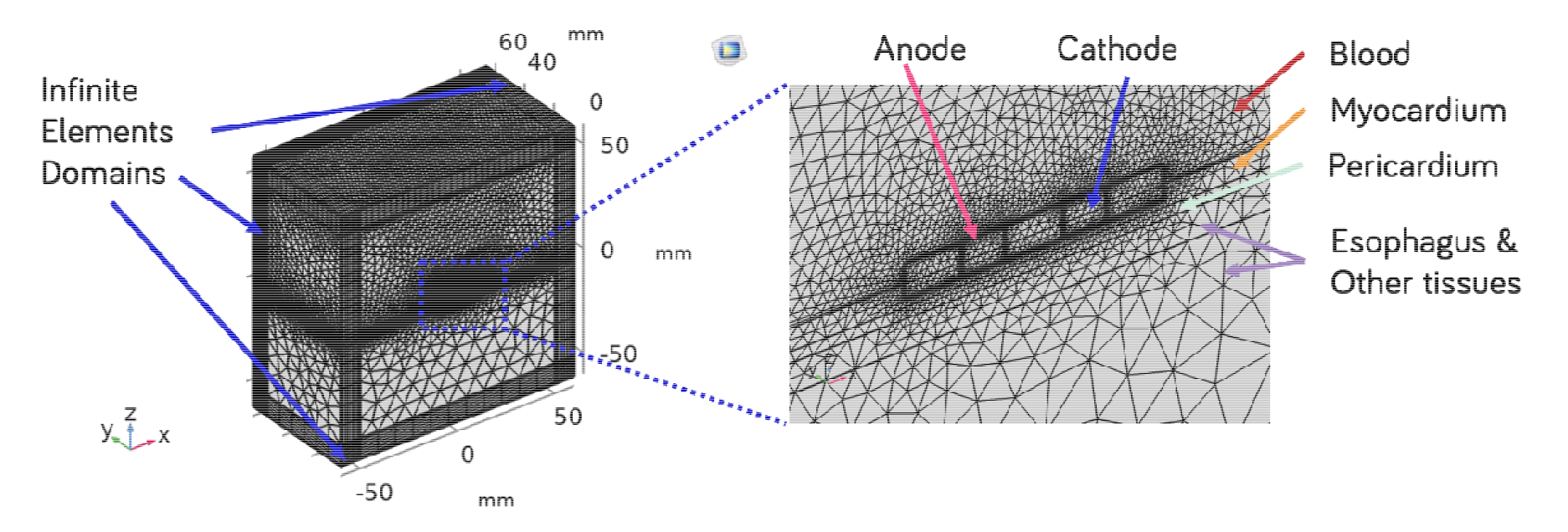
Convergence of geometry and mesh (446034 elements), featuring targeted refinements around the electrodes, ensuring enhanced resolution and accuracy in critical regions following successful convergence or independence criteria

### Solver independence

Finally, the solver independence also shows convergent results as presented in **Figure 7**. The default solver for fully coupled approach crashed (case 2), so the related bars are missing in the graphs. All the measured relative errors 1-3, 3-4 and 1-4 fulfilled the independence criteria. Accordingly, we chose the better solver based on a balance between small relative error and less computational time. In general, the fully coupled approach in 3D is considerably more computationally demanding than segregated approach. Accordingly, the case 4, segregated improved solver was chosen as the one fulfilling the independence criteria.

**Figure 7.**
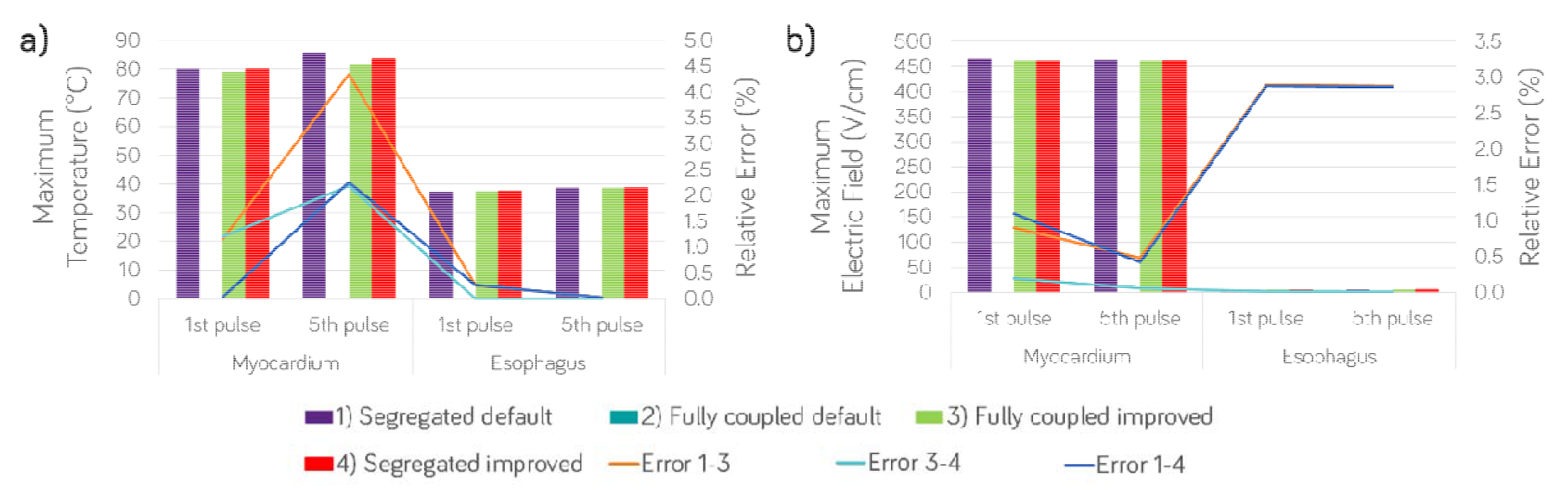
Solver independence for 1.5 kV, comparing a) maximum temperature and b) maximum electric field in both the myocardium and esophagus at the end of the first and fifth pulse for four different solving approaches. The case 2) did not converge, so the relative errors for temperature and electric field between case 1 and 2, 2 and 3, 1 and 3 are presented as lines and read on the right y-axis.

## Results

### Evolution of current, power and energy calculations

The evolution with time of current and power during the application of PFA pulses with peak voltage valued of 1000 V, 1500 V and 2000 V are presented in **Figure 8**. It is seen how the current increases with time for all the applied voltages. This is expected, as the electroporated cells are causing the tissue impedance to fall due to the creation of pores in the cell membrane. For 1000 V and 1500 V, the current and power increases with and exponential tendency, reaching a steady state and not considerably increasing more. Similarly, for 2000 V, it seems an exponential, but the value continues to increase slightly, suggesting the increase in voltage is non-linearly affecting the electroporated volume.

**Figure 8.**
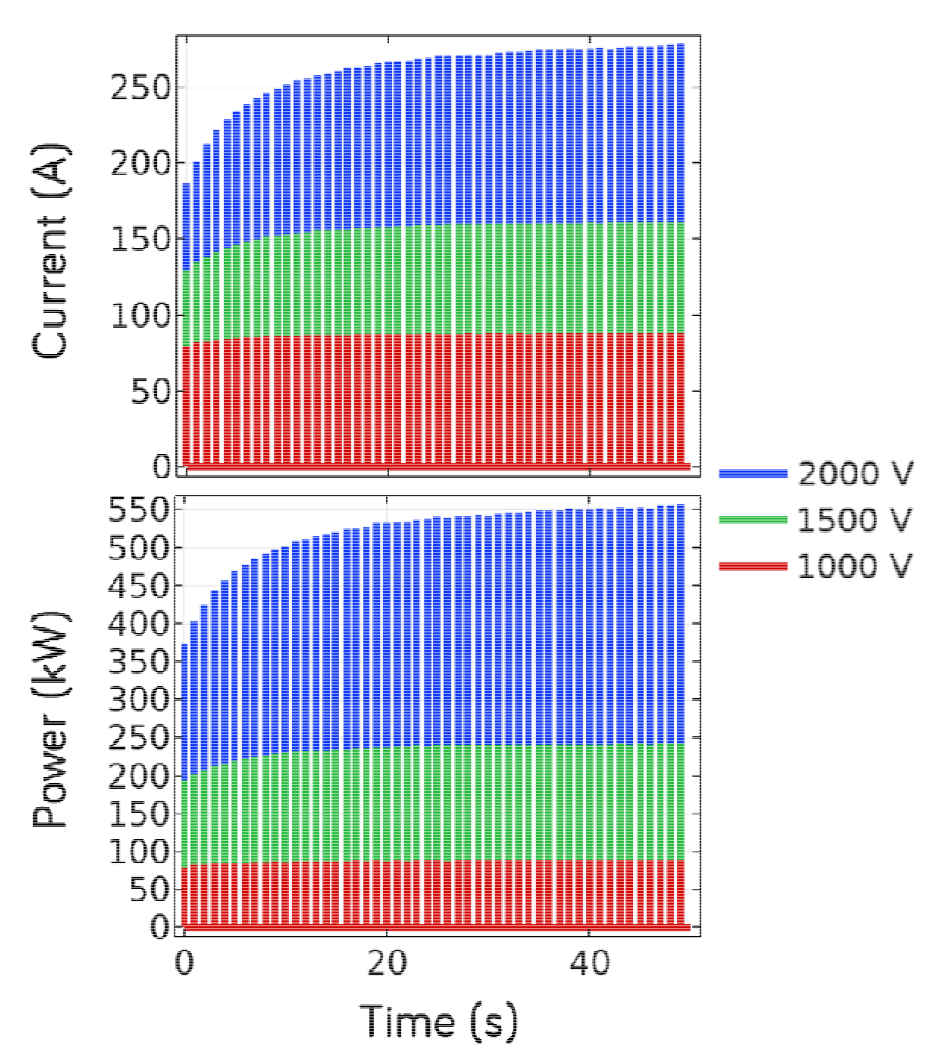
The evolution with time of current (top) and power (bottom) for 50 monophasic pulses.

From the power integral over time, the total energy (for the 50 pulses) is calculated and based on this value over the 50 applied pulses, the average energy per pulse is also calculated for the three peak voltages studied. Total energy for 1kV, 1.5kV, and 2kV pulse peak voltages, are 214 J, 570 J, and 1282 J, respectively. Average energy per pulse for 1kV, 1.5kV, and 2kV pulse peak voltages, are 4.28 J, 11.4 J, and 26.7 J, respectively.

### Electroporated tissue

Figure 9. presents the distribution of electric field strength for the three peak voltages considered in this study. The electric field inside the electrodes is zero as they are considered as perfect conductors. The myocardium, fat layer and esophagus, are respectively denoted as “myo”, “fat” and “eso”. As expected, the electroporated area is extended mainly into the cardiac tissue, and not considerably in the esophagus. This makes sense with both the clinical evidence of lower probability of the esophagus being electroporated, and the values of and for the esophagus in the electrical conductivity definition (Equation 9). The white contours, corresponding to the esophagus IRE threshold (1500 V/cm), are not entering the esophageal area, supporting again that there is no numerical evidence from this study, suggesting any esophageal electroporation.

### Thermal effects of PFA in the esophagus

Despite the fact that these results suggest that the esophagus is not electroporated during PFA (at least with the considered protocol), the thermal effects are in contrast not negligible. As presented in **Figure 10**, the temperature profiles for the three peak voltages considered at the end of the 50^th^ pulse suggest that 1000 V peak voltage would cause some heat in the esophagus, but not a significant amount. On the other hand, using 1500 V results in the maximum esophageal temperature close to the lethal isotherm of 50º C, and using 2000 V results in temperatures over 60º C.

**Figure 9.**
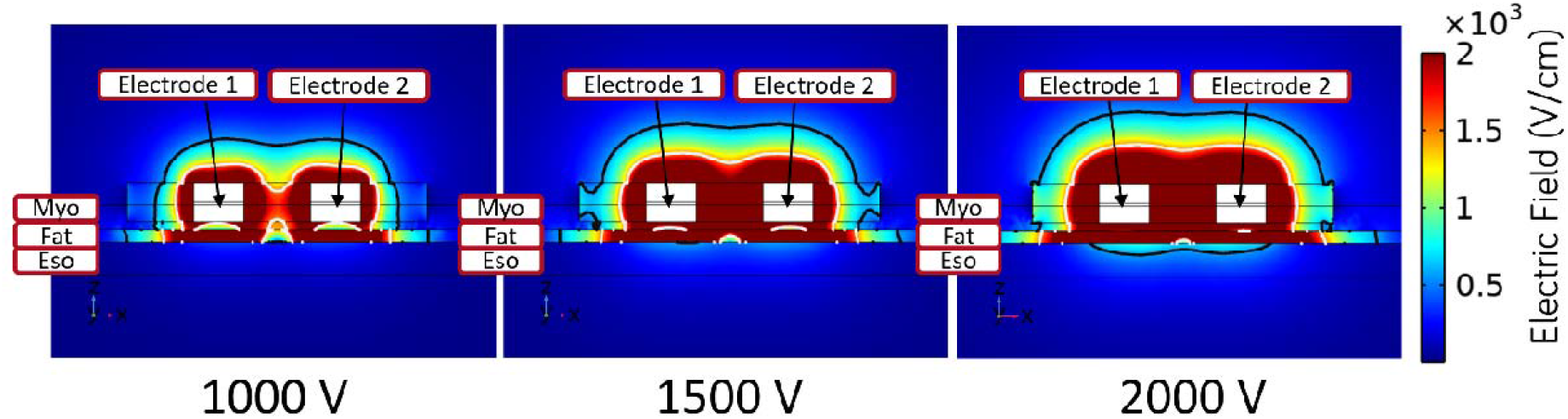
Electric Field in the x-y symmetry plane for different pulse peak voltages. Black contours are for 580 V/cm (Myocardium IRE Threshold). White contours are for 1500 V/cm (Esophageal IRE Threshold)

**Figure 10.**
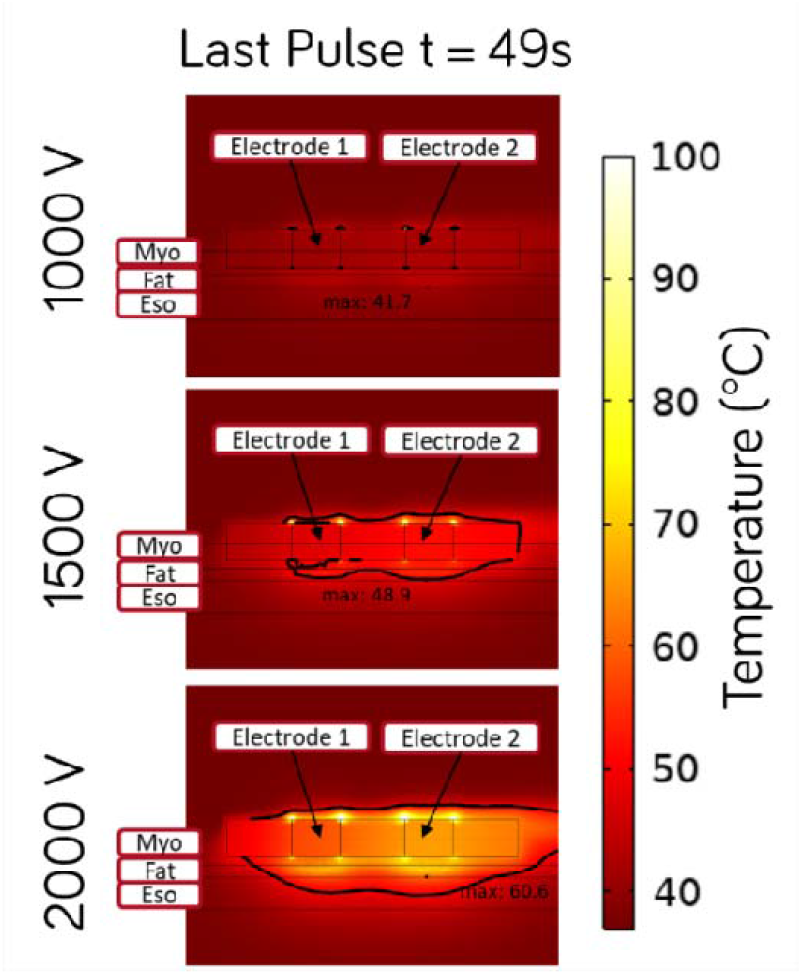
Temperature profile and maximum temperature in the esophagus at the end of the last pulse (t = 49s). The black contour corresponds to the 50 ºC lethal isotherm in the tissues and the blood.

The accumulated thermal effect is most pronounced at 1500 V and 2000 V, with an apparent non-linearly increasing behavior on the thermal damage by increasing the pulse peak voltages. This behavior has been also seen by other authors in Aycock et al.[18], Faroja et al.[5], and van Gemert et al.[4], supporting the reliability of the numerical results.

## Discussion

The in-silico model presented here offers a high-quality and efficient modeling approach to further investigate the thermal effects that occur with PFA before undertaking preclinical or clinical studies. Data continue to emerge demonstrating the thermal effects occurring from PFA, such that an in-silico model can serve as a method to more efficiently evaluate various operating parameters that may mitigate thermal effects and the adverse outcomes that can result.[7, 8] Kirstein et al. found that median esophageal temperature change was statistically significant and increased by 0.8□±□0.6°C, p□<□.001, with an esophageal temperature increase□≥□1°C observed in 10/43 (23%) patients, and the highest esophageal temperature of 40.3°C.[7] Measurement of the temperature profile of focal point, monopolar biphasic PFA on perfused thigh muscle of swine found that maximum average temperature rise for PFA was 7.6 °C, 2.8 °C, and 0.9 °C at the surface, 3-mm depth, and 7-mm depth, respectively.[8]

The model described here agrees well with data published in the field of oncology, where pulsed field ablation of tumors has been used for over a decade. Faroja et al. measured tissue temperatures in a porcine model using a commercially available system (Nanoknife Tissue Ablation System, Angiodynamics).[5] Using voltage settings of up to 3kV and pulse durations of 100 microseconds, they found post ablation temperatures ranging between 34° C for low-voltage 1500 V applications and 90 pulses to 84° C for high-voltage 2900V applications and 360 pulses. All combinations of voltage greater than 2500 V for 90 or more pulses produced temperatures greater than 50° C, which were associated with gross and histopathologic findings of thermal coagulation.[5] Dunki-Jacobs et al., and Agnass et al. subsequently reported similar findings of considerable heating sufficient to cause thermal damage, while also noting an exacerbating influence from the presence of metal within the ablation field, [6, 19] Mathematical models of oncologic applications have also reported significant thermal effects.[4] Thermal effects were more likely to occur for higher voltages (≥2000 V) and higher number of electrodes, but high-quality studies are needed to improve the predictability of the combined effect of variation in parameter combinations.[20] Nevertheless, in the field of oncology, data showing the thermal effects from PFA are abundant, with fistulas (typically vesico-cutaneous) occurring in up to 10.6% to 20% of patients.[5, 6, 19-22] As such, the importance of awareness of thermal effects from PFA cannot be overstated, and the availability of a model to further investigate these effects is valuable.

## Conclusions

Thermal effects from PFA occur and can be quantified with in-silico modeling. The model described here offers an efficient means of determining temperature effects from PFA over various operating conditions. Energy levels used clinically appear to have the potential to induce collateral thermal injury with repeated applications of pulsed field energy.

## Acknowledgements & Funding

This study was funded by the Spanish Ministerio de Ciencia e Innovación, Agencia Estatal de Investigación, Fondo Europeo de Desarrollo Regional (Grants PID2022-136273OA-C33 funded by MICIU / AEI/ 10.13039/501100011033 and by ERDF/EU, and the RYC2022-036965-I funded by MICIU/AEI/10.13039/501100011033 and ESF+), and the National Heart, Lung, And Blood Institute of the National Institutes of Health under Award Number R44HL158375 (the content is solely the responsibility of the authors and does not necessarily represent the official views of the National Institutes of Health).

## Notes

### Competing Interest Statement

Financial support for this research comes from consultancy provided by IN SILICO STEM to Attune Medical. Additionally, the software license for simulation was provided by IN SILICO STEM.
E. K. Employment with Haemonetics
We confirm that all relevant financial relationships have been disclosed and that they have not influenced the content of this manuscript.

